# Predicting P-glycoprotein Substrate Status Using a Pretrained Graph Neural Network: A TDC Benchmark Study

**DOI:** 10.64898/2026.06.01.729343

**Authors:** Jingjing Yan, Weicong Duan

**Affiliations:** QuantisMol LLC, Orange County, California, USA; College of Computing, Georgia Institute of Technology, Atlanta, Georgia, USA

**Keywords:** P-glycoprotein, ADMET prediction, graph neural network, deep learning, drug discovery, molecular property prediction

## Abstract

P-glycoprotein (Pgp/ABCB1) is a critical efflux transporter that significantly impacts drug bioavailability and multidrug resistance. Accurate prediction of Pgp substrate status is essential for early-stage drug discovery. In this study, we evaluate a pretrained Graph Isomorphism Network (GIN) with attribute masking on the Pgp_Broccatelli benchmark from the Therapeutics Data Commons (TDC). Our approach fine-tunes a GIN encoder pretrained on approximately 2 million molecules using a self-supervised attribute masking strategy, followed by a multilayer perceptron (MLP) classification head. On the TDC benchmark, our model achieves an AUROC of 0.937 ± 0.004 across five independent runs, ranking second on the leaderboard, as of May 2026. We further compare this approach against an XGBoost baseline using Morgan fingerprints (AUROC 0.912 ± 0.007), demonstrating the advantage of graph-based molecular representations with transfer learning for small-dataset ADMET prediction tasks.

## 1 Introduction

P-glycoprotein (Pgp), encoded by the ABCB1 gene, is a membrane-bound efflux transporter expressed in intestinal epithelium, blood-brain barrier endothelium, hepatocytes, and renal proximal tubule cells (Giacomini et al., 2010). As an ATP-dependent drug efflux pump, Pgp actively transports a broad range of structurally diverse compounds out of cells, thereby reducing intracellular drug concentrations. This efflux activity has two major implications for drug development: (1) reduced oral bioavailability when Pgp pumps drugs back into the intestinal lumen before absorption, and (2) contribution to multidrug resistance in cancer cells that overexpress Pgp (Szakács et al., 2006).

Given the clinical significance of Pgp-mediated efflux, computational prediction of Pgp substrate status has become an essential component of in silico ADMET (Absorption, Distribution, Metabolism, Excretion, and Toxicity) profiling. The Therapeutics Data Commons (TDC) provides a standardized benchmark for evaluating machine learning models on ADMET prediction tasks, including the Pgp_Broccatelli dataset comprising 1,218 molecules with experimentally determined Pgp substrate labels (Huang et al., 2021 Huang et al., 2022).

Recent advances in graph neural networks (GNNs) have shown promise for molecular property prediction by operating directly on molecular graph structures, where atoms are nodes and bonds are edges (Gilmer et al., 2017). A key innovation has been self-supervised pretraining of GNNs on large unlabeled molecular datasets, enabling effective transfer learning to downstream tasks with limited labeled data (Hu et al., 2020). Among pretraining strategies, attribute masking, which trains the model to predict randomly masked node and edge attributes, has proven particularly effective for learning transferable molecular representations.

In this work, we apply a pretrained GIN with attribute masking to the Pgp_Broccatelli benchmark and evaluate its performance against traditional fingerprint-based approaches.

## 2 Methods

### 2.1 Dataset

The Pgp_Broccatelli dataset (Broccatelli et al., 2011), accessed through the TDC Python API (PyTDC v1.1.15), contains 1,218 molecules labeled as Pgp substrates (Y=1) or non-substrates (Y=0). The dataset is split using the TDC’s official scaffold split: 973 molecules for training/validation and 245 molecules for testing. The training/validation set is further divided into 851 training and 122 validation molecules using TDC’s `get_train_valid_split` function with five different random seeds (1–5) to ensure robust evaluation.

### 2.2 Molecular Representation

#### 2.2.1 Graph Representation (GNN approach)

Each molecule is represented as a graph *G* = (*V, E*), where *V* is the set of atoms (nodes) and *E* is the set of bonds (edges). Node features encode atomic properties including element type, degree, formal charge, hybridization state, aromaticity, and hydrogen count. This representation preserves the full topological structure of the molecule without information loss from hashing.

#### 2.2.2 Fingerprint Representation (Baseline)

For comparison, we also represent molecules using Morgan circular fingerprints (Rogers and Hahn, 2010) with radius 2 and 2,048 bits, computed using RDKit. Each bit in the fingerprint vector indicates the presence or absence of a particular molecular substructure pattern within a two-bond radius of each atom.

### 2.3 Model Architecture

#### 2.3.1 Pretrained GIN with Attribute Masking

Our primary model uses a 5-layer Graph Isomorphism Network (GIN) encoder pretrained with attribute masking (Hu et al., 2020). The pretraining was performed on approximately 2 million molecules from ZINC15 and ChEMBL using a self-supervised objective: random attributes of nodes (atoms) and edges (bonds) are masked, and the model is trained to predict the masked attributes from the surrounding molecular context.

The pretrained GIN encoder produces a fixed-dimensional molecular embedding, which is passed to a classification head consisting of a two-layer MLP with hidden dimensions [512, 128]. The entire model, both the pretrained encoder and the classification head, is fine-tuned end-to-end on the Pgp_Broccatelli training set.

The model is implemented using DeepPurpose (Huang et al., 2021b) with DGL (Wang et al., 2019) and DGLLife backends. Training configuration: 100 epochs, learning rate 0.0005, batch size 128, Adam optimizer, binary cross-entropy loss.

#### 2.3.2 XGBoost Baseline

As a baseline, we train an XGBoost gradient-boosted classifier (Chen and Guestrin, 2016) on 2,048-bit Morgan fingerprints. Hyperparameters are optimized via grid search over max_depth ∈ {4, 6, 8}, learning_rate ∈ {0.01, 0.05, 0.1}, and n_estimators ∈ {300, 500, 800}, with early stopping (patience=50) on the validation set.

### 2.4 Evaluation

Model performance is evaluated using the area under the receiver operating characteristic curve (AUROC) on the held-out test set of 245 molecules. Following TDC guidelines, we report the mean and standard deviation across five independent runs with different random seeds for training/validation splitting, ensuring that the test set remains fixed.

## 3 Results

### 3.1 AttrMasking GNN Performance

Table 1 summarizes the per-seed performance of our AttrMasking GNN model.

**Table 1:** Per-seed test set performance of AttrMasking GNN on Pgp_Broccatelli.

| Seed | AUROC | AUPRC | F1 |
| --- | --- | --- | --- |
| 1 | 0.943 | 0.951 | 0.876 |
| 2 | 0.930 | 0.946 | 0.856 |
| 3 | 0.937 | 0.946 | 0.859 |
| 4 | 0.937 | 0.944 | 0.881 |
| 5 | 0.939 | 0.947 | 0.861 |
| **Mean ± Std** | **0.937 ± 0.004** | **0.947 ± 0.002** | **0.867 ± 0.010** |

**Table 2:** Comparison of AttrMasking GNN vs. XGBoost baseline.

| Model | Molecular Representation | AUROC | Parameters |
| --- | --- | --- | --- |
| AttrMasking GNN | Molecular graph | $0.937 \pm 0.004$ | 2,067,053 |
| XGBoost | Morgan fingerprint (2048-bit) | $0.912 \pm 0.007$ | N/A |

**Table 3:**
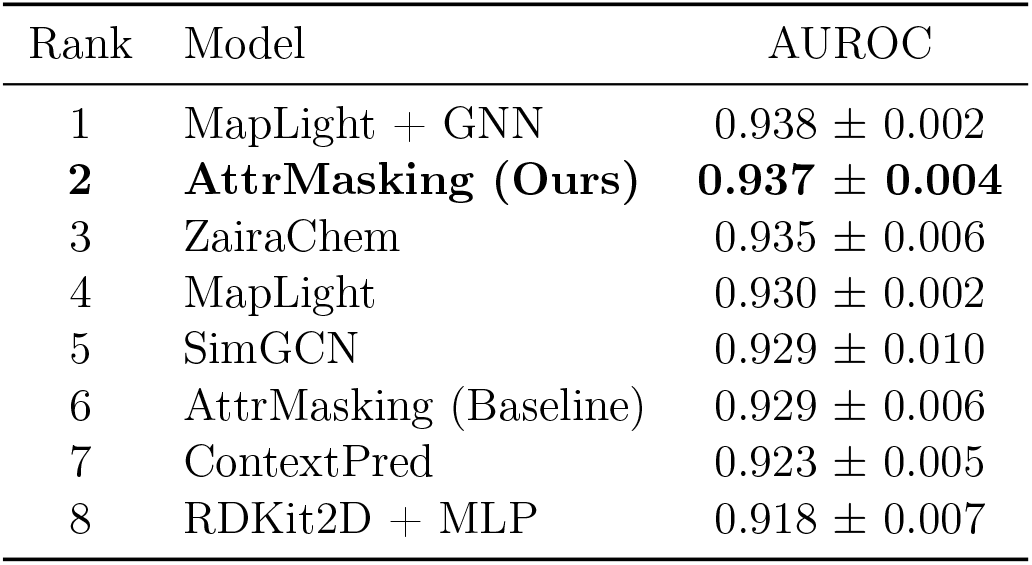
TDC Pgp_Broccatelli leaderboard (top entries).

| Rank | Model | AUROC |
| --- | --- | --- |
| 1 | MapLight + GNN | $0.938 \pm 0.002$ |
| 2 | **AttrMasking (Ours)** | **$0.937 \pm 0.004$** |
| 3 | ZairaChem | $0.935 \pm 0.006$ |
| 4 | MapLight | $0.930 \pm 0.002$ |
| 5 | SimGCN | $0.929 \pm 0.010$ |
| 6 | AttrMasking (Baseline) | $0.929 \pm 0.006$ |
| 7 | ContextPred | $0.923 \pm 0.005$ |
| 8 | RDKit2D + MLP | $0.918 \pm 0.007$ |

The model achieves consistent performance across all five seeds, with a standard deviation of only 0.004 in AUROC, indicating robust predictions regardless of the training/validation split.

### 3.2 Comparison with XGBoost Baseline

The AttrMasking GNN outperforms the XGBoost baseline by 2.5 percentage points in mean AUROC, demonstrating the benefit of graph-based representations with pretrained encoders over fixed molecular fingerprints.

### 3.3 Leaderboard Comparison

Our AttrMasking model achieves the second-highest AUROC on the leaderboard (0.937), only0.001 behind the leading MapLight + GNN model (0.938)(as of May 2026). Notably, our result improves upon the existing AttrMasking baseline (0.929) by 0.8 percentage points, likely attributable to differences in the classification head architecture and training hyperparameters.

### 3.4 Training Dynamics

Figure 3 shows the training loss and validation AUROC curves for Seed 1. The training loss decreases steadily over 100 epochs, from 0.698 to approximately 0.190. The validation AUROC rises rapidly in the first 10 epochs (from 0.927 to 0.949), then continues to improve more gradually, reaching a peak of approximately 0.963 around epoch 68 before stabilizing. The gap between validation AUROC (∼0.96) and test AUROC (0.943) suggests some degree of distribution shift between the validation and test splits, which is expected with scaffold-based splitting.

**Figure 1:**
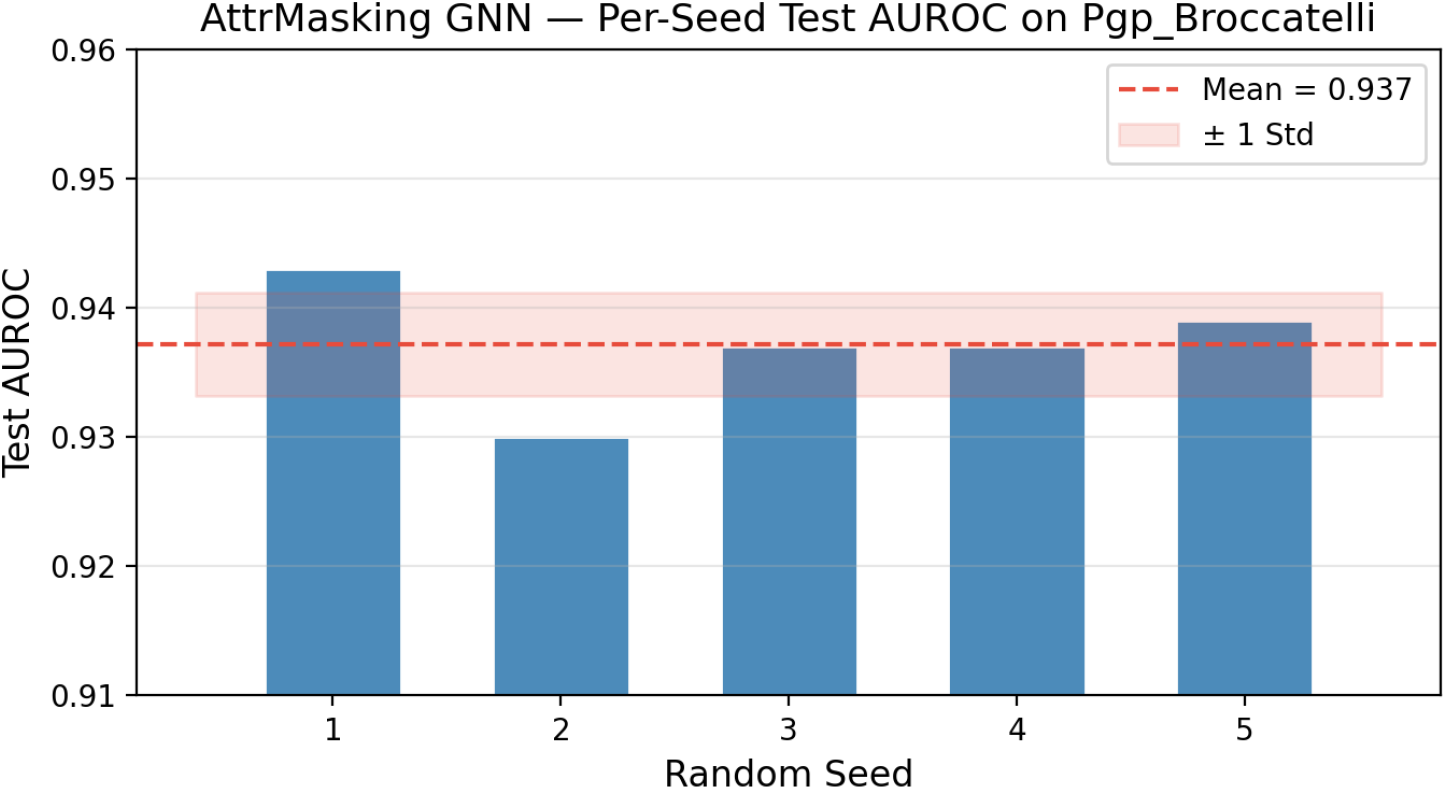
Per-seed test AUROC of the AttrMasking GNN model on the Pgp_Broccatelli dataset. The red dashed line indicates the mean AUROC (0.937), and the shaded region represents ± 1 standard deviation (0.004). Performance is consistent across all five random seeds.

**Figure 2:**
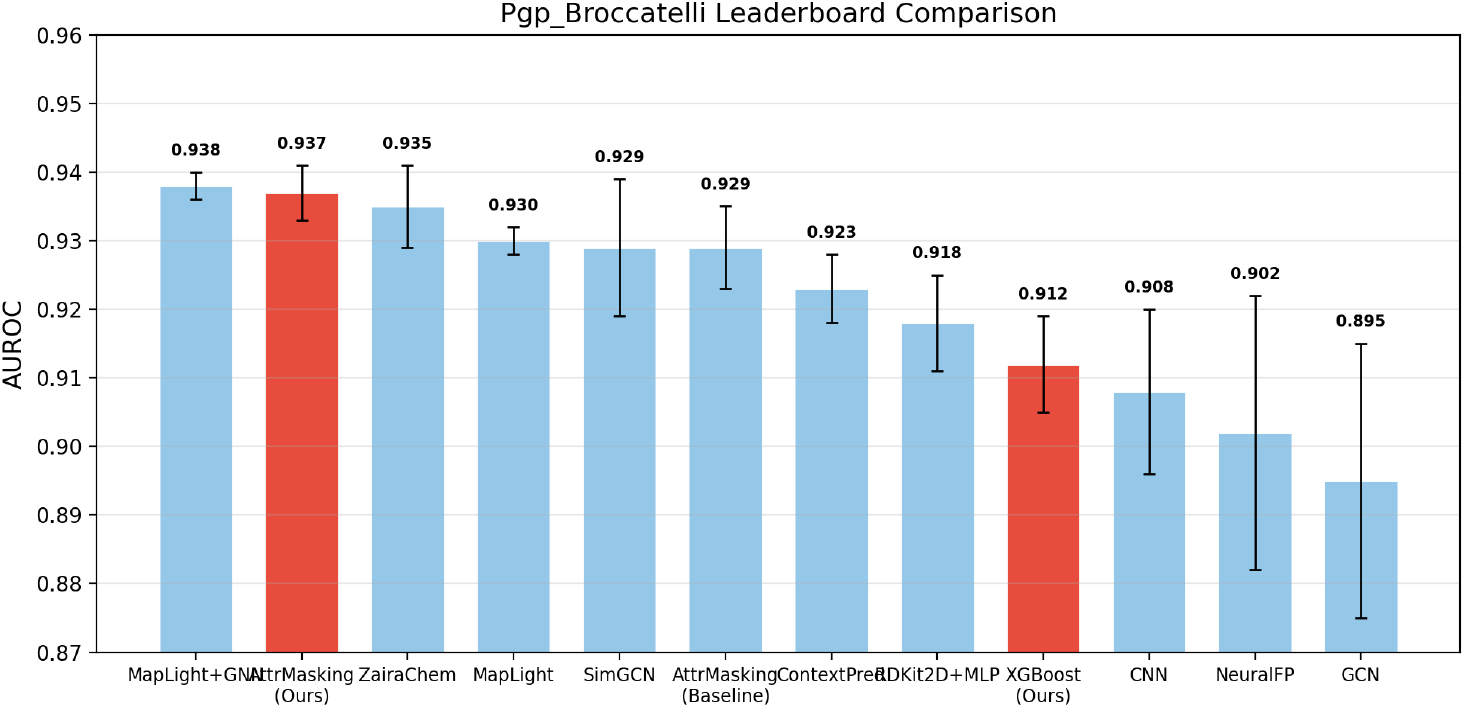
Comparison of model performance on the TDC Pgp_Broccatelli leaderboard. Red bars indicate our submissions (AttrMasking GNN and XGBoost baseline). Error bars represent standard deviation across five independent runs.

**Figure 3:**
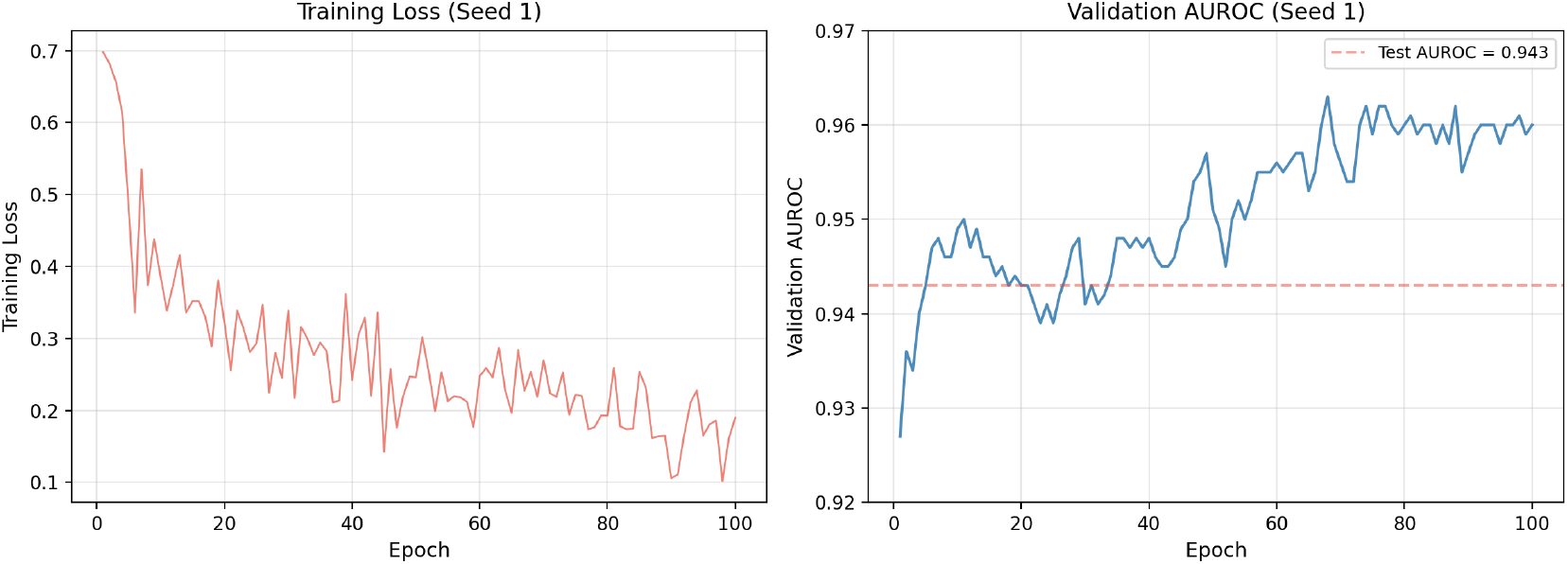
Training dynamics for Seed 1. Left: training loss per epoch showing steady convergence. Right: validation AUROC per epoch, with the final test AUROC (0.943) indicated by the red dashed line. The validation AUROC stabilizes around epoch 60–70.

## 4 Discussion

### 4.1 Pretrained GNNs vs. Fingerprint-Based Models

Our results confirm the advantage of pretrained graph neural networks over traditional fingerprint-based approaches for molecular property prediction on small datasets. The XGBoost model with Morgan fingerprints, despite being a strong baseline (AUROC 0.912), is limited by the information loss inherent in the hashing process: distinct substructures may map to the same fingerprint bit (collision), and spatial/stereochemical information is not captured by default.

In contrast, the GIN operates directly on the molecular graph, preserving full structural information. More importantly, the attribute masking pretraining enables the model to learn generalizable molecular representations from 2 million unlabeled molecules, effectively transferring this knowledge to the downstream Pgp prediction task despite having only 851 labeled training examples. This transfer learning paradigm is particularly valuable in drug discovery, where labeled data is often scarce and expensive to obtain.

### 4.2 Comparison with Existing AttrMasking Results

Our AttrMasking model (0.937) substantially outperforms the existing AttrMasking entry on the TDC leaderboard (0.929). Both use the same pretrained GIN encoder, so the performance difference stems from the fine-tuning configuration. Our model uses a larger classification head ([512, 128] vs. the default), a lower learning rate (0.0005), and trains for 100 epochs. These findings suggest that the fine-tuning strategy, not just the pretrained encoder, significantly impacts downstream performance.

### 4.3 Limitations

Several limitations should be noted. First, the Pgp_Broccatelli dataset contains only 1,218 molecules, which limits the statistical power of our evaluation. Second, our model does not account for stereochemistry: enantiomers with different Pgp substrate profiles would receive identical graph representations unless chiral features are explicitly included. Third, the scaffold split, while more realistic than random splitting, may not fully capture the challenges of predicting Pgp substrate status for truly novel chemical scaffolds encountered in drug discovery campaigns.

### 4.4 Future Directions

Several avenues for improvement exist: (1) incorporating stereochemical information (chirality and cis/trans isomerism) into node and edge features, (2) ensembling the GNN predictions with fingerprint-based models to combine complementary information sources, (3) exploring other pretraining strategies such as contrastive learning, and (4) extending this approach to multi-task learning across all 22 TDC ADMET endpoints simultaneously.

## 5 Conclusion

We demonstrate that a pretrained GIN with attribute masking achieves state-of-the-art performance (AUROC 0.937 ± 0.004) on the TDC Pgp_Broccatelli benchmark, ranking second on the leaderboard. The success of this approach highlights the value of self-supervised pretraining and transfer learning for molecular property prediction, particularly in the data-scarce regime common to ADMET profiling. Our code and results are publicly available at https://github.com/jingjingyan1/TDC-ADMET-Pgp-AttrMasking.

